# Control of electromechanical waves in engineered tissue of human iPSC-cardiomyocytes using a Halbach array and magnetic nanoparticles

**DOI:** 10.1101/2024.11.01.621542

**Authors:** Maria R. Pozo, Yuli W. Heinson, Christianne J. Chua, Emilia Entcheva

## Abstract

The Halbach array, originally developed for particle accelerators, is a compact arrangement of permanent magnets to create well-defined magnetic fields without heating. Here, we demonstrate its use for modulating the speed of electromechanical waves in cardiac syncytia of human stem cell-derived cardiomyocytes. At 40-50 mT magnetic field strength, a cylindrical dipolar Halbach array boosted the conduction velocity, CV, of excitation in a directional manner by up to 25% when the magnetic field was co-aligned with the electromechanical wave (but not when perpendicular to it). To observe the effects, a short-term incubation of the cardiac cell constructs with non-targeted magnetic nanoparticles, mNPs, was sufficient. This increased CV anisotropy, and the effects were most pronounced at slower pacing rates. Instantaneous formation and re-arrangement of elongated mNP clusters upon magnetic field rotation was seen, thus creating dynamic structural anisotropy that may have contributed to the directional CV effects. This approach may be useful for anti-arrhythmic control of cardiac waves.

**One sentence summary:** A Halbach array of permanent magnets can modulate the speed of excitation waves in human cardiac cell assemblies with magnetic nanoparticles.

## Introduction

The interest in deploying various energy sources for precise actuation and control of biological tissues, with and without genetic targeting of specific areas, has grown in the last decade(*1-4*). Light, ultrasound and magnetic fields are considered, among others. In addition to their clinical use for imaging (in magnetic resonance imaging, MRI), magnetic fields have been an attractive tool to explore for control of biological function, as for example in transcranial magnetic stimulation, because of their ability to penetrate deeper in the tissue, compared to light.

The Halbach array(*5*) is an arrangement of permanent magnets that can create well-controlled relatively strong magnetic fields at a smaller form-factor compared to those generated by electromagnets, without heating and at strengths that are bio-compatible. Originally, it was developed at the Lawrence Berkeley National lab in the 1980s to focus particle accelerator beams; recently, it has been deployed in the electric Tesla cars as part of the brushless motor.

In biomedical applications, the Halbach array’s compactness has been appreciated for building portable low-cost low strength (<100mT) MRI as a point of care devices(*6-8*). Other biomedical applications over the last decade include enhanced drug delivery(*9, 10*), magnetically assisted separation of biological substrates in suspension(*11-13*), and aggregation of different size particles(*14*). Of special interest, the recent work by Blümler and colleagues showcases different designs of cylindrical Halbach arrays for control and guidance of particles and cells in suspension(*15-18*).

Inspired by this work, we wondered if a Halbach array can be used to influence and control higher-level functional properties of biological tissues. This study reports on the use of a Halbach array to modulate the behavior of excitation waves in macroscopic syncytia of human induced stem cell derived cardiomyocytes, iPSC-CMs, mimicking human cardiac tissue in vitro. We explored different orientations of the applied magnetic field with respect to the direction of cardiac waves and quantified the speed of such waves when the cardiac syncytia were placed in a Halbach array without and with short-term administration of magnetic nanoparticles. The work has translational implications for potential magnetic control of cardiac arrhythmias.

## Results

### The Halbach array provides a well-defined and strong magnetic field at a compact size and simple design, compatible with cell culture incubators and imaging setups

A cylindrical Halbach array (design and photos in **Figs. 1A-B**) and an array holder were constructed to facilitate control and imaging of cardiomyocyte monolayer samples in 35 mm dishes with glass bottom. Magnet polarity in the inner and outer rings (**Fig. 1C**) followed a design similar to (*15*) to achieve the desired fields (**Fig. 1F**). The visualized magnetic fields were as theoretically expected (**Fig. 1D**). The measured field strength generated by the inner ring alone (with magnets in a dipolar configuration) was approximately 44-47 mT and quite uniform. Variation in repeated measures with a handheld gaussmeter was approximately ± 1.5 mT, which we deemed acceptable. The magnetic field was fully contained within the Halbach array, as predicted. Considering the very compact formfactor and small magnets, this is an excellent field strength. Control of the direction of the magnetic field (zero magnetic force) was accomplished by rotating the inner ring (**Fig. 1D**, middle). The addition of a concentric outer ring (in a quadrupolar configuration) adds the possibility to control not only the direction but also the magnetic force. When rotating the outer ring, a predictable non-uniform field can be created (**Fig. 1D**, right). For example, the addition of the outer ring and its rotation at an angle β (approx. 15°), created a magnetic field gradient (34 mT to 52 mT).

**Fig 1.**
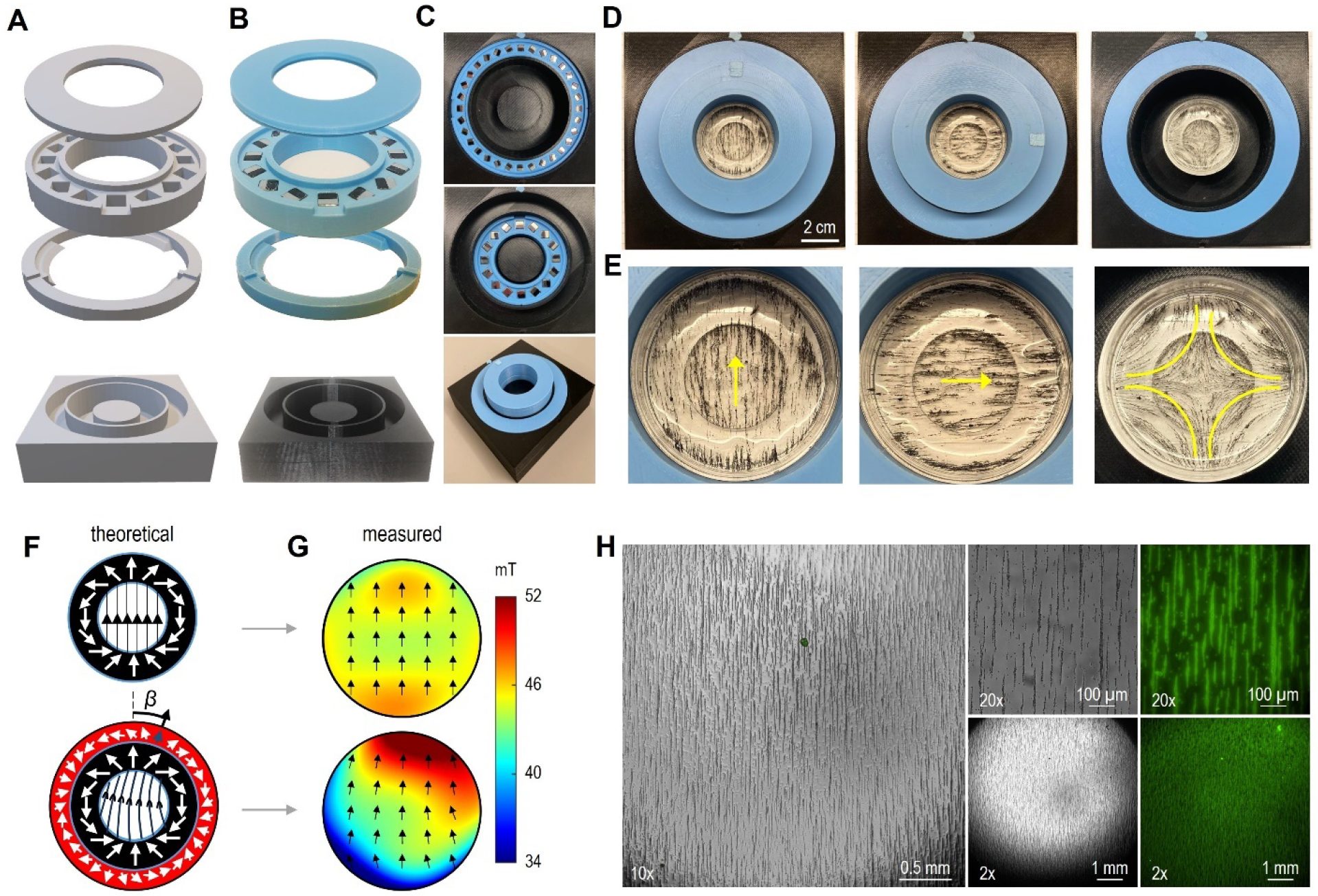
Design of a compact Halbach array for cell culture and visualization of the magnetic fields. **A-B**: CAD drawing and photos of the inner rings; **C**.Outer rings fit concentrically in holder component with freedom to rotate. **D-E**. Magnetic fields, visualized with iron shavings in viscous liquid for the inner ring with no rotation, inner ring with 90° rotation, and outer ring alone, with close-ups. **F**. Theoretical fields for inner ring alone (F, upper) and inner and outer rings together, with slight outer ring rotation at angle β≈15° (F, lower), white arrows indicate magnet polarity and black center arrows illustrate the corresponding magnetic field lines. **G**. Actual measurements for the configurations in F, black arrows indicate measured magnetic field direction; magnetic field strength (in mT) encoded by color. **H**. Oriented extracellular matrix by the Halbach array: magnetic nanoparticles (mNPs) were mixed with fluorescein-labelled fibronectin to coat cell culture dishes and positioned in a Halbach array in the incubator overnight; shown are brightfield and fluorescence images under different magnification.

To determine whether the presence of the magnetic nanoparticles (mNPs), combined with the application of a uniform magnetic field, could influence the structural organization of the extracellular matrix, we used fluorescently-labeled fibronectin mixed with mNPs. Placement in the Halbach array overnight (with only the inner ring) resulted in macroscopic alignment/anisotropy of the extracellular matrix (**Fig. 1H**). In comparison, control experiments using fibronectin with mNPs put on top of a magnetic plate instead of the Halbach array did not produce aligned samples.

### Cardiac conduction velocity can be boosted along the direction of the Halbach array-generated magnetic field (<50 mT) even shortly after administration of mNPs

Next, we set out to quantify the effects of the Halbach array-generated magnetic fields on the electromechanical function of engineered syncytia of human iPSC-CMs, without and with the addition of mNPs. To do so, we designed a hollow-bottom holder for the Halbach array and integrated it with an in-house built all-optical electrophysiology imaging system(*19*), **Fig. 2**. This system allowed optical mapping of voltage and calcium waves while the direction of the magnetic field was controlled with respect to such paced excitation waves. Only the inner ring of the Halbach array was used in these experiments. The experimental design is outlined in **Fig. 2C**, and it included multi-frequency pacing (0.5 Hz, 0.75Hz, 1Hz) at three defined magnetic field directions (α = 0°, 90°, 180°) with respect to the excitation wave, pre-and post-administration of mNPs. So, for each sample, at least 18 different records were obtained for voltage or calcium (9 before and 9 after the two-hour incubation with mNPs).

**Fig 2.**
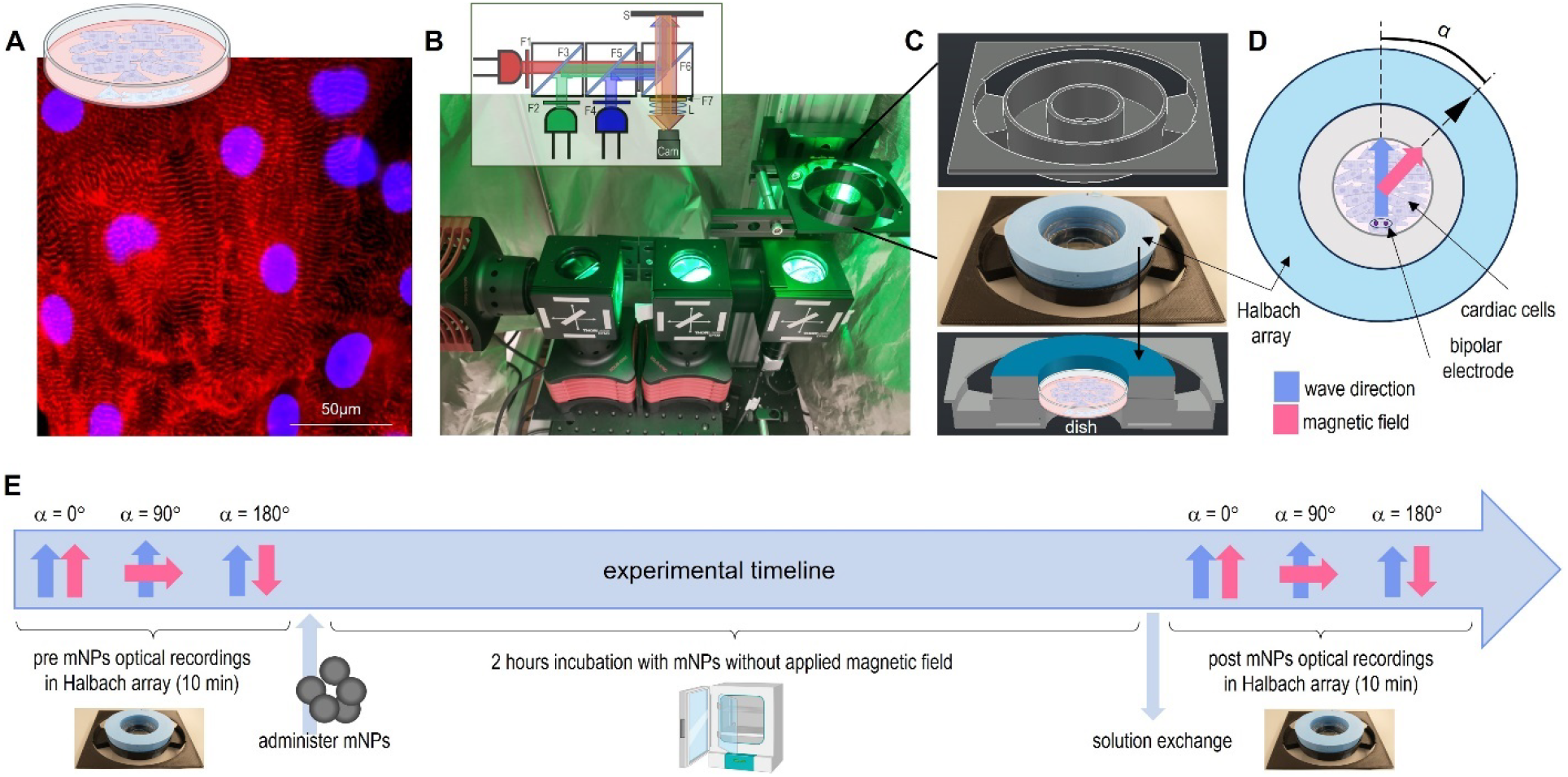
Integrated optical mapping system with a Halbach array holder for a 35mm dish along with the experimental timeline. **A**.Example image of hiPSC-CMs cultured in glass-bottom dishes, with labeled alpha-actinin and nuclei. **B**. OptoDyCE-plate imager for optogenetic pacing, and optical mapping of voltage and calcium, F1: 655/30; F2: 525/30; F3: 605LP dichroic; F4: 470/24; F5: 495LP dichroic; F6: 473/532/660 dichroic; F7: 595/40+700LP; L: camera lens; S: sample stage; Cam: CMOS camera. **C**. The custom designed holder-array-sample arrangement with hollow accessible bottom sits on the OptoDyCE stage. **D**. Within the array (top view) the bi-polar electrode is indicated, initiating an excitation wave (blue arrow), while the magnetic field direction (pink arrow) is changed by rotating the Halbach array, e.g. α = 0°, 90°, and 180°. **E**. Experimental timeline: optical recordings for three magnetic field directions pre-and post-mNP incubation (in most experiments 2 hours); at each field orientation the sample was paced at multiple frequencies and maps were analyzed. Biorender was used in part of the image.

Activation maps were constructed to quantify the effects of the applied magnetic fields on excitation wave propagation, **Fig. 3**. Samples exhibited expected conduction velocity (CV) restitution effects (CV slowing with faster pacing) without or with mNPs (**Figs. 4A-B**). An increase in CV (speed up) was noted, especially along the direction of the applied magnetic field, after incubation with mNPs for two hours. CV effects could be seen both in voltage and calcium wave imaging. This effect was quantified more robustly with multiple samples over different cultures and confirmed the increased CV after mNP incubation (**Fig. 4C**), especially when the magnetic field was along the same direction as the pacing wave (α = 0°, blue bars). At this orientation, the effect was frequency dependent -CVs increased by 25.8% at 0.5 Hz pacing, 23.7% at 0.75 Hz pacing, and 16% at 1.0 Hz pacing. In most of our experiments, samples were imaged with α = 0° recorded first, followed by a Halbach array rotation to α = 90° and 180° and the statistically significant CV boost was only observed in the α = 0° orientation, with a trend of increase for the 180°C. We decided that the quick-succession measurements while rotating the Halbach array in the presence of mNPs may exhibit some residual torque effects. We decided to test an alternative order of the imaging (recorded α = 180° first, followed by α = 90° and 0°) and this resulted in a significant CV boost observed in the α = 180° orientation (**Fig. 4D**). Importantly, no CV increase was seen when α = 90° orientation was recorded first (**Fig. 4E**), confirming that alignment of the magnetic field, regardless of polarity orientation, is important for the observed boost in CV in the presence of mNPs. Also, no bulk temperature difference was seen in samples with and without mNPs in the Halbach array, so the observed CV boost cannot be explained by thermal effects.

**Fig 3.**
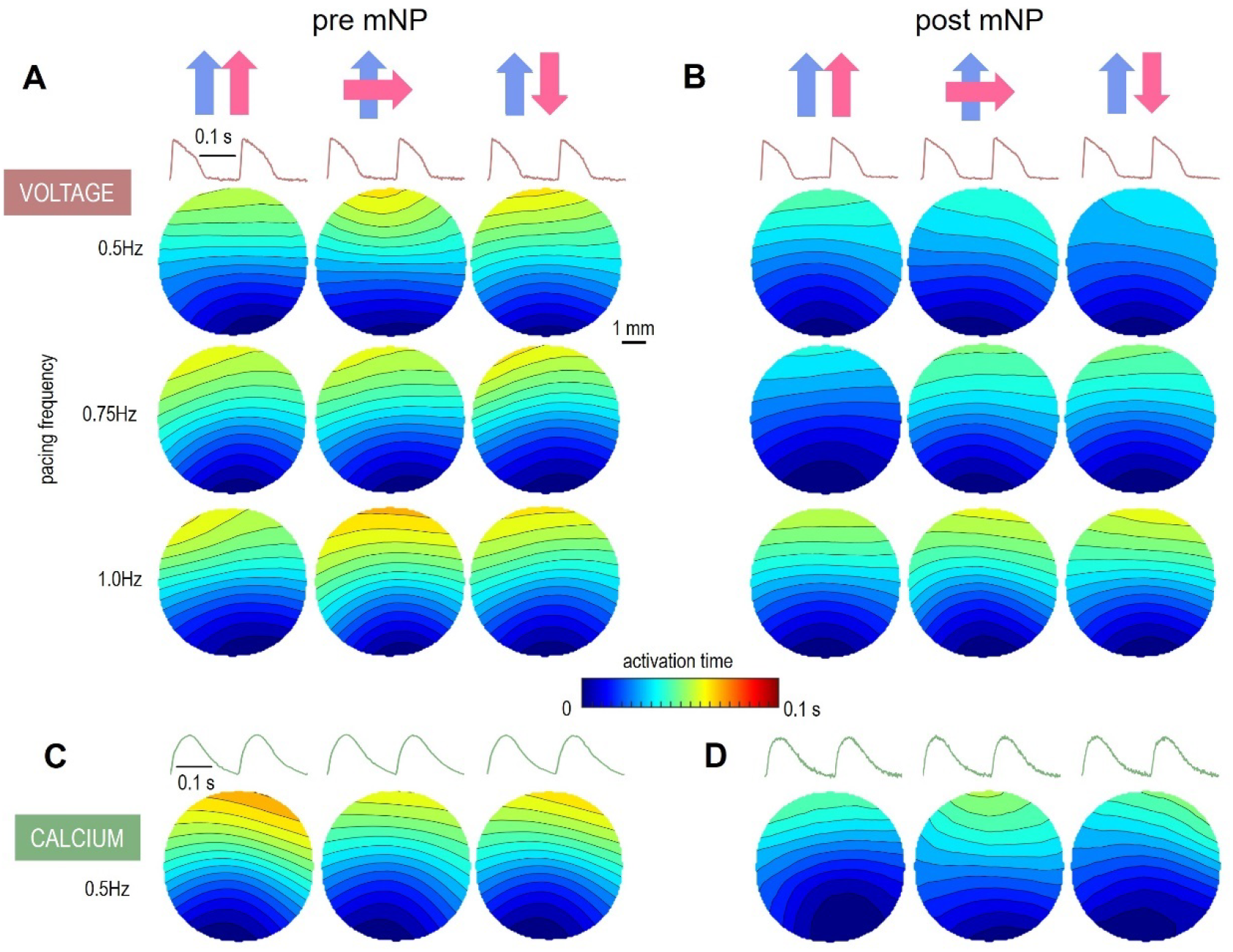
Activation maps of triggered cardiac waves (voltage and calcium) in static magnetic fields without and with mNPs. **A-B**.Voltage-based traces and activation maps for three pacing frequencies before (A) and after (B) two-hour incubation with mNPs. Blue arrows indicate the direction of the excitation wave and pink arrows indicate the magnetic field direction. The shown maps are from the same cardiomyocyte sample. **C-D**. Calcium-based traces and activation maps before (C) and after (D) mNP incubation. All maps in C and D represent the same cardiomyocyte sample. The scale bars indicate 1s and 1mm and the color bar indicates the wavefront activation times (in seconds).

**Fig 4.**
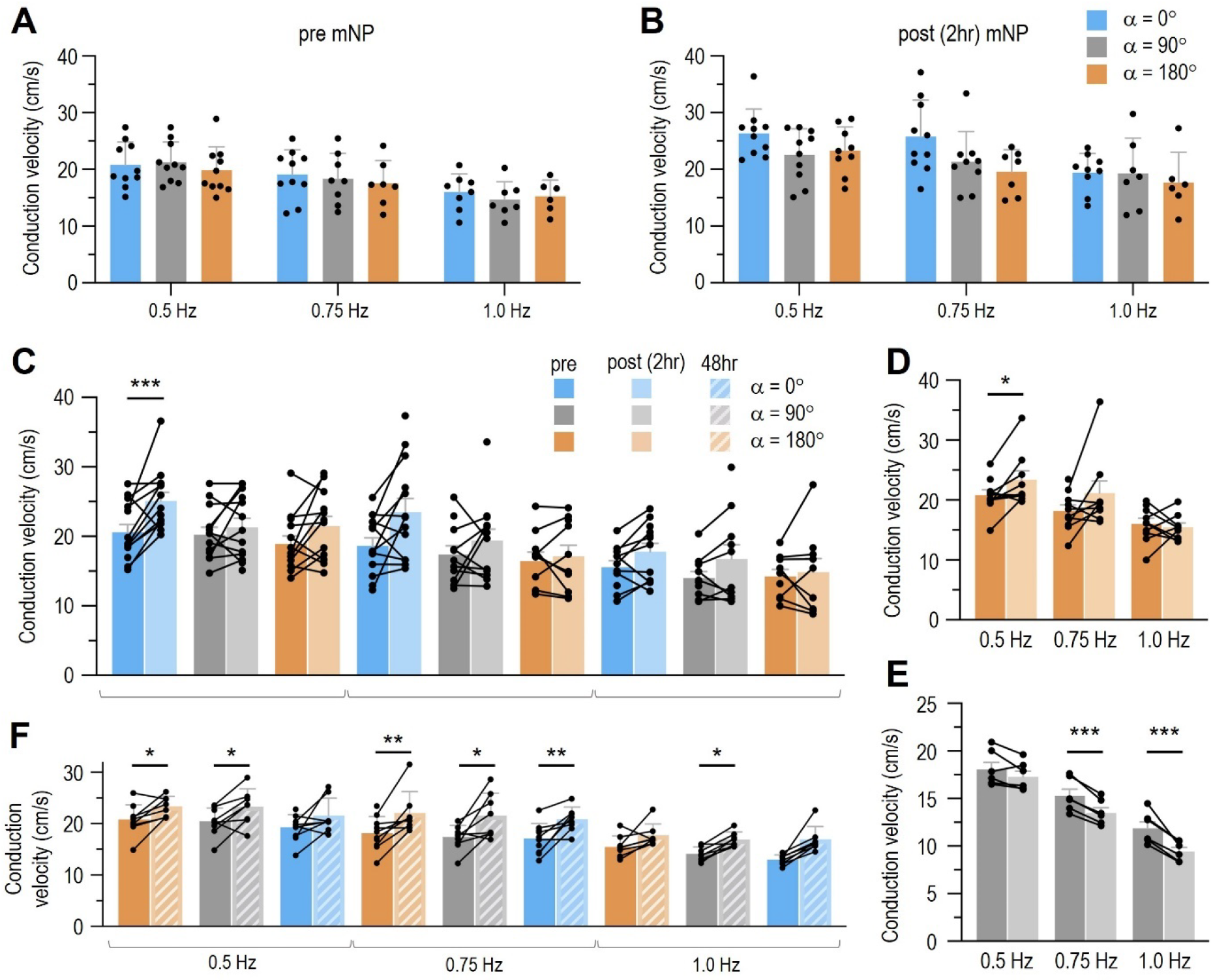
Quantification of the effects of static magnetic fields on cardiac wave conduction velocity (CV) pre and post 2hr and 48hr incubation with mNPs. **A-B**.CV (cm/s) in paced hiPSC-CM at α = 0°, 90°, and 180° is displayed for pre-mNP measurements (A) and after 2hr mNP incubation (B). **C**. Alternative display of CV in matched pairs before and after mNP incubation, grouped by pacing frequency and field rotation α; n=9-13 matched pairs. **D**. Alternative order of the experiments, CV in paced hiPSC-CM before and after mNP incubation where α = 180° was recorded first instead of last. **E**. CV in paced hiPSC-CM before and after mNP incubation where α = 90° was recorded first; n=6 matched pairs. **F**. Longer incubation with mNPs (48 hrs) and quantification of CV measured, with α = 180°, followed by 90° and 0°; n=10 matched pairs.*p<0.05, **p<0.01, ***p<0.005.

Observation of this acute effect raised questions about the longer-term effects of mNPs on CV. We saved samples from a typical mNP experiment and repeated the imaging protocol again 48 hrs later. Unlike the immediate effects, here increase in CV was seen across multiple array orientations and pacing frequencies (**Fig. 4F**). The cells were still healthy at this time point and in the week following these measurements, corroborating the biocompatibility of these mNPs without adverse cell viability effects with long term exposure.

### Magnetic field (<50mT) application alone or mNPs alone do not alter conduction velocity

Previous work on nerve cells has demonstrated that strong magnetic fields (around 1T, about 20 times stronger than used here) applied to isolated sciatic nerve may result in increased CV along the fiber, and that this effect can linger even after the magnetic field is removed for over 20 min(*20*), although other reports showed contradictory results(*21*). We conducted several control experiments to check if, at the much lower magnetic field strengths used here (< 50mT), the magnetic field exposure itself may alter CV, **Fig. 5A**). The resulting activation maps (examples in **Fig. 5B**) for these control experiments show no significant change in conduction velocity (**Fig. 5C**) two hours after initial magnetic field exposure, thus precluding direct effects on CV by the magnetic field at these strengths.

**Fig 5.**
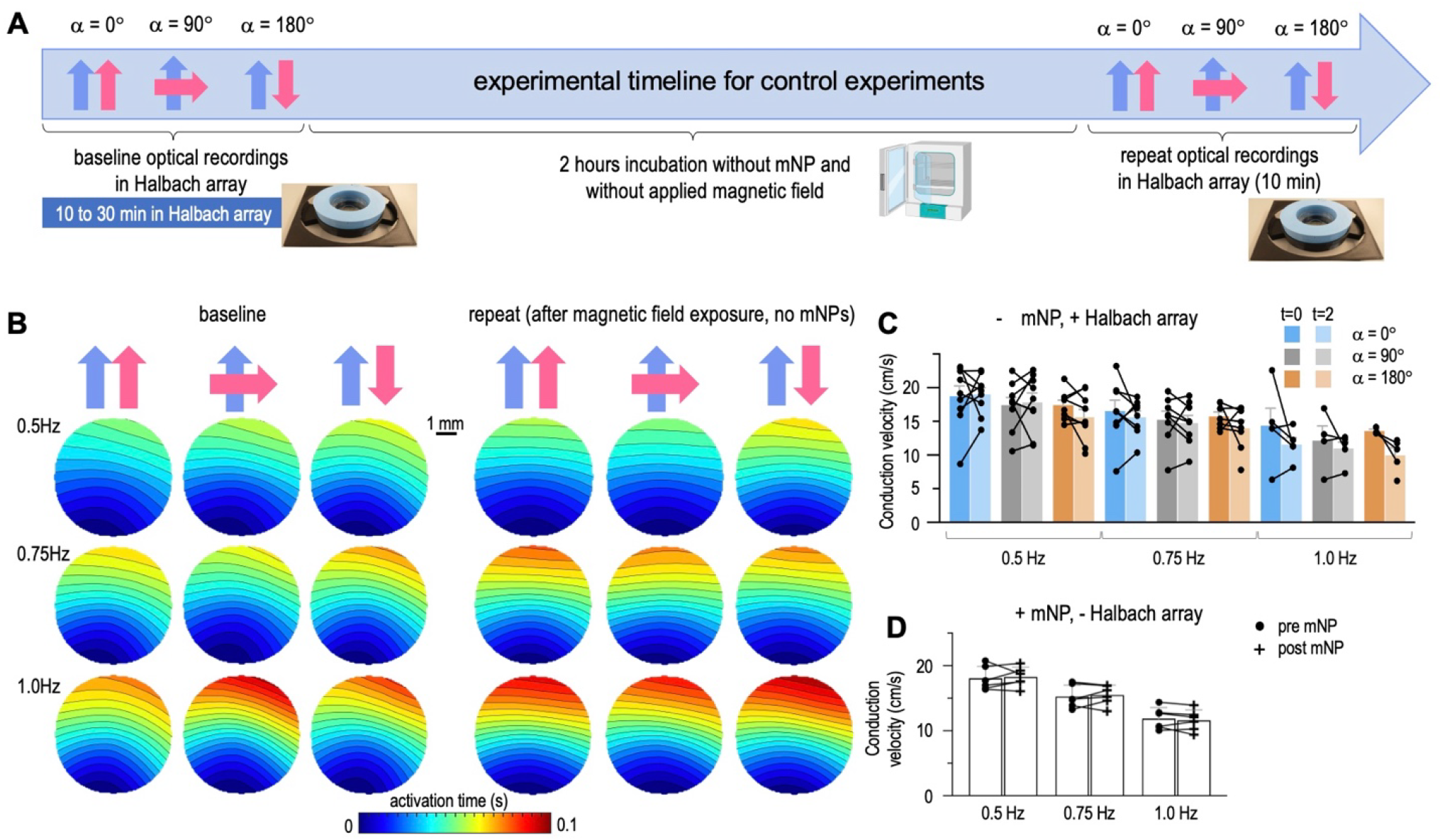
Control experiments for potential residual effects on CV of static magnetic fields without mNPs or with mNPs alone without applied magnetic fields. **A**.Experimental design and timeline. **B**. Voltage-based example activation maps at varying pacing frequencies and field directions, before and after 2hr incubation (no mNPs). The scale bar indicates 1mm and the color bar indicates wavefront activation times (in seconds). **C**. CVs from matched pairs are grouped by pacing frequency and field rotation α; n=3-9 matched pairs. **D**. CVs for hiPSC-CMs that were treated with mNP for 2hr and then imaged without the application of a magnetic field; n=6 matched pairs. Biorender was used in part of the image.

Furthermore, we wondered if the presence of the mNPs, without applied external magnetic field, may affect cardiac CV. Results from such control experiments are shown in **Fig. 5D**, with no changes in CV at the 2 hr time point post mNP administration. Therefore, in this study neither the magnetic field exposure alone nor the short-term mNP administration alone could have caused the observed changes in CV; both must be present to achieve CV increase.

### Conduction velocity increase could be explained in part by instantaneous mNP-mediated increase in structural anisotropy along the direction of the magnetic field

From the experiments conducted, we noted that the conduction velocity ratio (CV when the magnetic field is along the direction of the wave vs. CV when the magnetic field is applied across the direction of the wave) is increased (CV ratio > 1) only in the presence of mNPs (**Fig. 6A)** and not in control experiments without mNPs (**Fig. 6B**). This suggested certain invoked anisotropy properties in the presence of mNPs during applied magnetic field as a possible mechanism dictating the wave speed.

**Fig 6.**
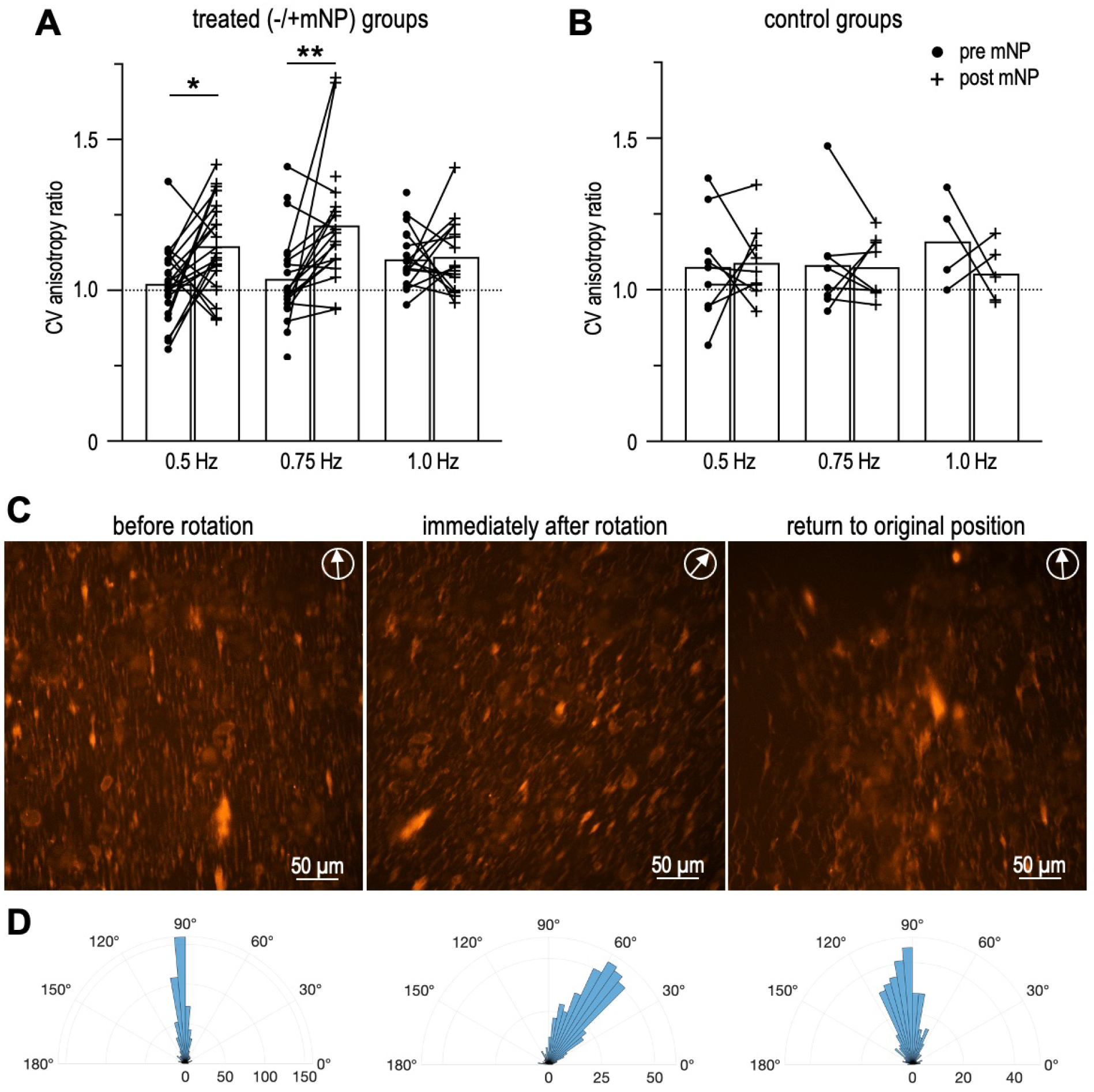
Magnetic field-induced anisotropy in conduction velocity. **A-B**.CV anisotropy ratios (CV with magnetic field along the wave / CV with the magnetic field perpendicular to the wave) in paced cardiomyocytes and field aligned with the direction of excitation; with mNPs n=23 matched pairs and without mNPs n=9 matched pairs. **C**. Visualization of fluorescently-labeled mNPs applied to cardiomyocytes under rotating magnetic field. **C**. Orientation of fluorescent magnetic microparticles was quantified at each array position, as indicated by the arrow in the upper right corner. **D**. Histograms of angles from the analyzed images, bottom numbers indicate the number of objects detected in the image used for orientation quantification. *p<0.05, **p<0.01.

We used fluorescently labelled mNPs to visualize the immediate responses to an applied magnetic field. As in other experiments, cardiomyocytes were incubated with the mNPs for 2 hr and solution was exchanged to remove free floating mNPs before imaging. Under the initially applied magnetic field from the Halbach array, the mNPs appeared to form clearly aligned clusters/structures (**Fig. 6C-D**, left). Rotation of the Halbach array immediately led to re-alignment of the assembled structures, although not as precisely as in the original configuration (**Fig. 6C**, middle). A rotation back to the original direction of the magnetic field introduced residual effects, and even though the general direction was restored immediately, the overall alignment was not as good. This is quantified with angular histograms which widened after rotations (**Fig. 6D)**. It should be noted that these fluorescently labeled mNPs were larger than the ones used in our functional mapping experiments (300 nm compared to 50 nm), as we did not have a fluorescently labeled version of the original small mNPs. In our experience, the smaller mNPs also tend to make anisotropic clusters, which are harder to visualize. We suspect that internalization of the mNPs into the cells introduces anisotropy during magnetic field application and the mNP clusters attempt to follow the direction of the field with immediate re-alignment, but the experienced torque may disrupt some of the clusters, especially bigger ones.

## Discussion

There has been a long-standing interest in magnetic fields (AC & DC, static, gradient, etc.) and how they may influence biological function, especially in clinically used modalities like MRI(*22, 23*). Some earlier studies have been concerned with potential adverse health effects of ambient and environmental magnetic field exposures(*24, 25*). Alternatively, recently, interest is growing in leveraging magnetic fields for precise control of biological processes. This can be done without augmenting tissues with magnetic actuators, as is the case for transcranial magnetic stimulation(*26*), or with the addition of magnetic materials in biomedical applications such as drug release, soft robotics, magnetic guidance, among others(*2*). The field of magnetogenetics, for example, in parallel developments with optogenetics(*1*), aims to deploy magnetic fields together with magnetic nanomaterials and genetically encoded targets for control of cellular processes, as reviewed in(*27-30*). To our knowledge, no prior studies have used a Halbach array to influence cardiac function or excitable tissue properties.

### Key findings and implications

In this study, we found that constant magnetic fields of modest strength (< 50mT) generated by a compact Halbach array can be used to modulate the conduction velocity of excitation waves in engineered cardiac tissue, after a short-term incubation with non-functionalized magnetic nanoparticles. The effects depend on the pacing frequency (more prominent at slower rates) and on the relative direction of the applied magnetic field with respect to the wave. About 25% increase in CV was observed when the magnetic field and the excitation waves were co-aligned. All cell experiments here were conducted with static magnetic fields and the Halbach configuration was with zero magnetic force. Concentric arrangements of a dipolar and quadrupolar Halbach arrays can further yield control over both the direction of the field and the magnetic force, and such an assembly can be computer controlled(*16*). Overall, our results are the first to show that such low magnetic fields, when combined with non-targeted mNPs can directionally modulate electromechanical waves. With proper control of the magnetic field strength and the magnetic force(*15, 16, 18*) and with optimized responsiveness to the magnetic force through mNP design, this approach can come close in performance to optogenetic “wave steering” of excitation(*31, 32*), with the added advantage of potentially being useful in deeper tissue access.

The speed (or conduction velocity) of excitation waves is an important property defining the dynamics and stability of neural and cardiac systems. Modulation of that key multicellular property provides a means of controlling the waves. No prior studies have found mT-strength magnetic fields to influence excitation properties, and we also did not detect any effects of the Halbach array on unmodified cardiac tissue. Older reports using much stronger magnetic fields (> 1T) on isolated nerves have yielded controversial results. Reno(*20*) reported directional boost of conduction velocity in sciatic nerve (when the magnetic field is aligned with the fiber but not when it is perpendicular) upon relatively short exposure to static magnetic fields and with lasting effects after field removal. Others have obtained contradictory results in similar configurations(*21*). With the addition of mNP, and with the much smaller field strengths of the Halbach array used here, we saw anisotropic CV boost, similar to (*20*).

### Possible mechanisms

Identifying the mechanism by which such modulation of excitation wave speed is possible at these low strength fields with a short-term application of mNPs is important for designing future control technology. However, this was outside of the scope of the current study and will require much more in-depth investigation. Much remains unknown about the interactions of magnetic fields with biological tissue, as exemplified with controversies from the magnetogenetics field(*33, 34*).

Conduction velocity of excitation waves is fundamentally most influenced by fast-activating depolarizing ion channels, by the conductance of inter-cellular connections (gap junctions and nanodomain-mediated electric coupling) as well as by the structural organization of the cells and matrix (anisotropy can boost CV). Without a genetic component in this study and with the immediate effects seen here, there are several possibilities for the observed results. Among the potential mechanisms are thermal effects mediated by the mNPs when in the Halbach array, mechanical effects of locally altered rigidity, tension, compression by the mNPs, instantaneous anisotropic mNP clustering and redirection of the ion current flow, locally experienced Lorentz forces. All of these can then engage classic intermediaries reported to be influenced by stronger magnetic fields or specifically targeted in magnetogenetics. These are mechano-sensitive ion channels and thermo-sensitive ion channels, endogenously expressed in the cardiomyocytes, activatable at relatively low threshold (forces less than 10 pN(*35, 36*)), and capable of generating depolarizing currents that can influence excitability and CV.

To this end, we have confirmed that within the Halbach array with or without incubation with mNPs, no measurable temperature differences were observed. This does not preclude very localized thermal gradients around the mNPs, which we could not capture with our thermal camera. However, at the low magnetic field strengths, such thermal effects are likely negligible, and even if present, would not explain the directionality of the effects on CV. In our study, we saw instantaneous anisotropic mNP cluster re-arrangements in response to the direction of the magnetic field, **Fig. 6**, which we believe to be important for the observed anisotropic CV effects. The nanoparticles used in our experiments are either endocytosed or attach to the cell membrane(*37*). When clusters of such attached mNPs are being re-oriented by the magnetic field, this can provide direct stimulation to various mechano-sensitive and other ion channels and cellular components. Further investigation will be needed to confirm such a hypothesis.

### Future developments and uses

This initial study shows a promise for further developments in the cardiac field. We focused on using one Halbach dipole cylinder to apply a constant magnetic field over the cardiac syncytia and we are excited about the possibility of applying dynamically changing fields by using both the dipole and quadrupole cylinders in tandem for more sophisticated wave control in the future, where the magnetic force is also controlled in addition to the field direction(*16*). The biocompatibility seen in our study is also a promising feature. The experiments did not involve substantial sample modification like magnetogenetics or optogenetics, apart from the two-hour incubation with mNPs, similar to those already FDA-approved as MRI contrast agents. While we have not yet rigorously investigated long-term viability of mNP-treated cardiomyocytes, we have tested effects up to 48 hrs after mNP treatment and saw healthy cells with high conduction velocities. With further studies confirming long-term compatibility, this opens possibilities for *in vivo* work leveraging the Halbach array design.

Specific uses in the cardiac field will benefit from the Halbach configuration and holder design described here, which are compatible with lab incubators and offer scalability with human iPSC-CM constructs. A further optimized version of the reported approach can include several important applications: 1) control of cardiac electromechanical waves as an anti-arrhythmic therapy *in vivo* and 2) speeding the integration of engineered cardiac tissue (out of human iPSC-CMs, preloaded with mNPs) for safe heart regeneration. Cardiac arrhythmias are often initiated by perturbed wave trajectories, as seen in myocardial infarction, for example. Current day anti-arrhythmic therapies (cardioversion and defibrillation) are brute-force approaches that do not involve gentle modulation of the aberrant waves, as proposed in “wave steering”(*31, 32*). With noninvasive label-free imaging of such wave disruptions(*38, 39*) and with magnetic guidance of the waves as suggested here, it may be possible to counter arrhythmias in new ways. With respect to the application to heart regeneration, interestingly, a recent study reported that the incorporation of mNPs improved integration of human stem-cell-derived cell grafts in rodent hearts by promoting better cell-cell contacts(*40*). The mechanisms of such improvement may be shared with some of the CV boost factors discussed here. If the new-design point-of-care MRIs based on Halbach arrays(*7, 8*) are further optimized to become suitable for cardiac use, such devices can be turned into all-magnetic systems (imaging + actuation), deployed for anti-arrhythmic interventions and aiding in cardiomyoplasty.

## Methods

### Halbach array design, assembly, and characterization

A cylindrical Halbach array was constructed with inner and outer rings within a holder to accommodate 35mm cell culture dishes. All parts were designed in AutoCAD® (v.2021, Autodesk, San Francisco, CA), **Fig. 1A-C**, similar to (*15*). Parts were printed using a Prusa i3 MK3S 3D printer and PLA plastic. Rings were filled with Magcraft® neodymium-iron-boron magnets (inner rings: 1/4” cubes, outer rings: 3/16” cubes, National Imports, Vienna, VA) arranged by polarity as shown, and press-fitted to assemble. Each holder accommodates two inner rings that are vertically stacked and one outer ring, which fits concentrically around the inner rings. Magnetic fields were visualized using iron shavings mixed with viscous fluid (uncured PDMS, Dow Silicones Corporation, Midland, MI). Measurements of magnetic field were done with a hand-held gaussmeter (HGM09s, MAGSYS, Dortmund, Germany) for inner ring stack alone and for inner ring stack with outer ring at slight angle β (**Fig. 1G**, upper and lower, respectively). Measured field values throughout the sample plane (one measurement at each black arrow in **Fig. 1G**) were linearly interpolated, followed by a 4^th^ degree polynomial fit to project the trend onto the larger circular area.

To integrate the Halbach array with our optical mapping system(*19, 41*), we redesigned the holder to have access beneath the sample, shown in **Fig. 2C**, to be able to optically image a 35mm glass-bottom cell culture dish from underneath, similar to the microscope use of such an array(*42*). The holder ensures the sample is concentric with the Halbach array while the Halbach array rotates within the holder around the sample (**Fig. 2C**, section plane view).

### Magnetic nanoparticle administration and visualization

For all functional cell experiments, we used 50 nm magnetic nanoparticles (mNPs), composed of Au-Fe2O3-Poly-L-Lysine (Nanoshuttle-PL, #657847, Greiner Bio-one, Kremsmünster, Austria). mNPs were administered to cells at 50 μL per dish, supplemented with 250 μL warmed opti-MEM or maintenance media (Fujifilm CDI, Madison, WI), and incubated at 37C for two hours or as indicated.

For visualization of the effects of the magnetic field on extracellular matrix protein assembly (**Fig. 1H**), micron-sized mNPs were used (Lenti-X™ Accelerator, #631257, Takara Bio, Kusatsu, Shiga, Japan). These particles were mixed with fluorescein-labelled fibronectin (488 nm, #FNR02, Cytoskeleton, Denver, CO), 50 μg/mL in PBS. Magnetic particles and fluorescent fibronectin were mixed in a 1:40 v/v ratio and applied to the glass portions of 35 mm dishes with 14 mm glass bottoms for 12 hours. Imaging was done using an inverted fluorescence microscope Nikon Ti-E.

For visualization of the immediate anisotropy-inducing effects of magnetic fields on particles and cells (**Fig. 6C**), fluorescently-labelled mNPs, 300 nm size (nanomag®-CLD-F, micromod Partikeltechnologie GmbH, Rostock, Germany) were administered to cells at 20 μL per dish, supplemented with 280 μL warmed maintenance media, and incubated for two hours. In all experiments, the solution was exchanged to remove free-floating magnetic particles prior to imaging using an inverted fluorescence microscope Nikon Ti-E at 20x magnification.

Temperature changes in sample solution were measured over time using a thermal camera (FLIR One Pro, Teledyne FLIR, Wilsonville, OR). Samples with and without mNP incubation (1mL bulk volume) were measured immediately after removal from incubator and measured again every minute for 14-16 minutes (approximating the length of time that samples spend outside the incubator during imaging protocol).

### Human iPSC-CM culture

Commercially differentiated human iPSC-derived cardiomyocytes (iCell Cardiomyocytes^2^, Fujifilm CDI, Madison, WI) were thawed and plated according to manufacturer’s protocol at 270,000 cells/dish to create dense syncytia. Prior to thawing, 35 mm dishes with 14 mm glass bottoms were coated overnight with fibronectin (#356008, Discover Labware Inc, Bedford, MA) at 50 μg/mL. Cells were grown in a 5% CO_2_ incubator at 37°C and fresh maintenance media was supplied to the cells every 48 hrs until 7 days post-thawing when measurements were conducted.

### Immunocytochemistry

Immunocytochemistry was used to visualize hiPSC-CM structure, **Fig. 2A**. Cardiomyocyte sarcomeres were labelled with monoclonal anti-α-actinin antibody (Catalog #A7811, Millipore Sigma) and nuclei were labelled with Hoechst (Catalog # H3570, Thermo Fisher Scientific). Cells were rinsed with PBS, fixed with 10% formalin, and permeabilized with 0.2% Triton X-100 in 5% FBS. Two-stage antibody labelling was applied and the samples were imaged with a Nikon Ti2 60x objective in oil.

### Experimental Design

On day 7 post-thaw, cells were fluorescently labeled for voltage and calcium detection, placed into the Halbach array holder with inner ring stack only (no outer ring for cell experiments) and placed onto the OptoDyCE-plate imager(*19*). A bipolar platinum electrode was used to initiate excitation waves in a specified direction from the edge of the sample. The Halbach array inner ring was rotated with respect to this direction of the wave, starting with α = 0° (blue and pink arrows, **Fig. 2D**). Paced voltage and calcium fluorescence signals (using 3 pacing frequencies) were collected (further described in *Optical setup*) at α = 0°, 90°, and 180°, through rotation of the inner ring with respect to the sample. Cell samples were then removed from the Halbach array and incubated with magnetic nanoparticles for two hours (**Fig. 2E**). Sometimes, cells were re-labeled for voltage and calcium to ensure strong signal. Then, samples were placed in the Halbach array and imaging/pacing protocol was repeated for α = 0°, 90°, and 180°. For some experiments, the order of imaging was reversed so that α = 180° was recorded first, followed by α = 90° and α = 0°. These samples were then maintained for another 48 hrs before the imaging/pacing protocol was repeated for a second time. In one experiment, only α = 90° was recorded before and after 2 hr mNP incubation. For no-mNP control experiment, timeline and imaging were replicated, but two-hour incubation was carried out with fresh, plain maintenance media (**Fig. 5A**). For no-array control experiment, timeline and two-hour mNP incubation were replicated but follow-up imaging was performed with no array in the array holder.

#### Dye labeling

To obtain voltage and calcium signals, day 7 post-thaw cells were labeled as described previously (*43*) with voltage dye BeRST1 at 1-2 μM (from Evan W. Miller, University of California, Berkeley) and/or calcium dye Rhod-4 AM at 10 μM (AAT Bioquest, Sunnyvale, CA) in warmed (37°C) opti-MEM media. Solution was replaced with 1 mL fresh, warmed opti-MEM or maintenance media before imaging on the in-house developed imaging system.

#### Optical setup

The OptoDyCE-plate imager(*19*) was used to collect optical recordings. It is a stand-alone low-cost high throughput all-optical system, previously used for parallel recordings from multiple samples in a 96-well or 384-well plate (photo and schematic in **Fig. 2B**). The high-power red and green LEDs are for excitation of voltage and calcium-sensitive dyes, respectively. Optical components information is available in(*19*). Using epi-illumination and the combination of various filters, voltage and calcium fluorescence signals were recorded with a low-cost camera, Basler acA720-520um (Basler AG, Ahrensburg, Germany), operating at 100 frames per second. Note the sample-Halbach array together have the same elevation level as the sample stage when the holder is placed on the sample stage, which ensures the image spatial resolution to be 132μm/pixel, as in (*19*).

#### Pacing protocol

Electrical stimulation was achieved by a bipolar platinum electrode, connected to a MyoPacer stimulator (IonOptix, Westwood, MA). Optical recordings were performed under 0.5, 0.75, and 1.0 Hz electrical pacing frequencies.

### Data/Image processing

A custom software adapted from(*44*) in MATLAB (v.R2023a, MathWorks, Natick, MA) was used to analyze the optical mapping data and obtain activation maps. A central region of interest was selected (11.9 mm diameter) to analyze the recordings. Locally weighted spatial Bartlett filter was applied with window size of 7 pixels and temporal filtering was applied with span of 0.2 s. From the activation maps, conduction velocity was calculated with detected activation times (selection of points along the wave). Anisotropy in conduction velocity was calculated as the ratio of CV along vs. across the direction of the wave.

The orientation of fluorescent microparticles under magnetic field was quantified using the regionprops function in MATLAB (v. 2023a, MathWorks, Natick, MA). Fluorescent images were binarized at uniform thresholds and the orientation values were filtered to remove very small detected objects (MajorAxisLength < 4) before plotting the angle histograms.

### Statistical Analysis and Reproducibility

Experiments were repeated over multiple cell cultures for each condition to confirm reproducibility. Statistical analysis was performed in GraphPad Prism (v.10.1.1, GraphPad Software, Boston, MA). Differences between matched pairs (before vs after mNP incubation or control) were identified by parametric paired t-tests. Multiple tests correction was not used here due to the very small number of tests.

## Author contributions

EE designed and oversaw the study, MRP conducted experiments, analyzed data, and organized all figures. CJC designed, printed, and assembled the original Halbach array assembly; MRP and YWH designed and printed the array holder and built additional Halbach arrays; YWH designed the imaging system and helped with many of the experiments and some of the analysis. MRP and EE wrote the manuscript with input from YWH.

## Acknowledgements

We acknowledge Julie L. Han and Morgan Pettebone who assisted several experiments and Grant Kowalik for technical assistance in the 3D printing process.

